# Direct Amide Bond Formation By A Spore Peptidoglycan Biosynthetic Enzyme

**DOI:** 10.1101/2025.03.28.644612

**Authors:** Madison E. Hopkins, Zachary Yasinov, Mark J. Fakler, Grace E. Wilde, Katherine G. Wetmore, Michael A. Welsh

**Affiliations:** Chemistry Department, Hamilton College, Clinton, New York, 13323, United States

## Abstract

The cortex layer of the peptidoglycan cell wall surrounding bacterial spores contains a modified sugar, muramic-δ-lactam, that is essential for spore germination. Genetic evidence has linked the conserved enzyme SwsB to the muramic-δ-lactam biosynthetic pathway. SwsB belongs to a large family of metal-dependent deacetylases, but its function is unclear because a putative catalytic residue is mutated. We have used native cortex peptidoglycan substrates to show that SwsB acts not as a deacetylase but as a monofunctional muramic-δ-lactam cyclase, the first enzyme reported with this activity. SwsB is remarkable in that it catalyzes lactam synthesis by direct intramolecular condensation of a carboxylate and primary amine with no apparent requirement for chemical energy input. SwsB will accept a minimal peptidoglycan substrate and, surprisingly, does not require a metal ion cofactor for cyclase activity. Our results suggest an *in vivo* role for SwsB and lay the foundation for mechanistic and structural studies of an unusual enzyme.

---

Enzymes that synthesize the peptidoglycan cell wall surrounding bacterial cells are established antibiotic targets.^1^ The biochemical steps of peptidoglycan biosynthesis in vegetative (i.e., growing) bacterial cells are well-characterized,^2^ but less attention has been paid to the pathway in bacterial spores, dormant cells that form in response to nutrient starvation.^3,4^ The processes of sporulation, in which a spore is formed, and germination, in which a spore reinitiates growth, each involve dramatic structural remodeling of peptidoglycan.^3^ Enzymes operating in these pathways are potential targets for spore-specific antimicrobials,^5,6^ but many of them remain poorly characterized.

Peptidoglycan is a carbohydrate polymer of alternating *N*-acetylglucosamine (GlcNAc) and *N*-acetylmuramic acid (MurNAc) residues (Figure 1a). Each MurNAc is substituted with a pentapeptide that crosslinks to other polymer strands. In the cortex layer of peptidoglycan surrounding mature spores, 30-50% of MurNAc residues are converted to muramic-δ-lactam (Figure 1a).^8–10^ In the model organism *Bacillus subtilis*, two enzymes are sufficient for muramic-δ-lactam synthesis.^11–13^ The L-alanine amidase CwlD first cleaves the pentapeptide from Mur-NAc. Next, the transamidase PdaA catalyzes hydrolysis of the MurNAc acetyl group and cyclization of the product, muramic acid (MurN), to give muramic-δ-lactam (Figure 1b).^12^ The muramic-δ-lactam modification is unique to spores and plays an essential role in their physiology. During germination, the entire cortex must be degraded by glycosidases, principally SleB and CwlJ, and muramic-δ-lactam residues serve as a molecular recognition motif to target these enzymes to the cortex layer.^14^ Spores without muramic-δ-lactam therefore fail to germinate.^15^

**Figure 1.**
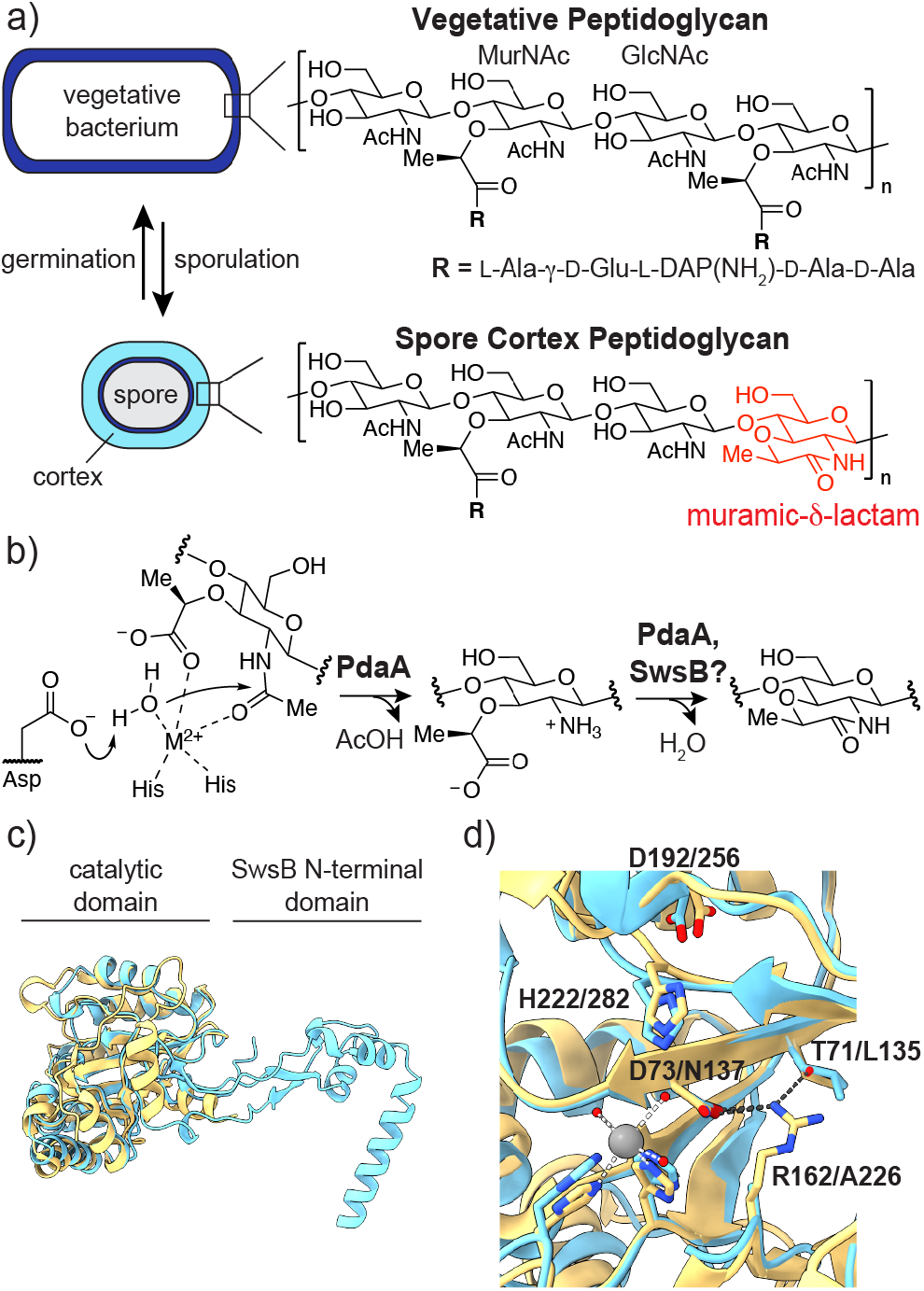
Spore cortex peptidoglycan contains muramic-δ-lactam. (a) Peptidoglycan structure in vegetative *B. subtilis* and the spore cortex. (b) Synthesis of muramic-δ-lactam by PdaA. MurNAc deacetylation is followed by MurN cyclization. (c) Aligned structures and (d) active site residues of *Bs*PdaA (yellow, PDB 1W1B^**7**^) and *Bs*SwsB (cyan, AlphaFold3 model). Labels indicate residue numbers in *Bs*PdaA and *Bs*SwsB, respectively. *Bs*SwsB residues 228-231 are hidden for clarity.

A recent genetic screen in *B. subtilis* revealed that an uncharacterized protein, SwsB, was required for CwlJ-dependent spore germination.^16^ SwsB is conserved in sporulating organisms^16,17^ and closely resembles PdaA (Figures S1, S2). Both are members of a family of polysaccharide deacetylases classified as carbohydrate esterase 4 (CE4) whose active site contains a divalent metal cation, typically Zn^2+^, liganded by two conserved histidine residues.^7,18,19^ The metal acts as a Lewis acid to enhance the nucleophilicity of a water molecule during deacetylation (Figure 1b).^20^ The transamidase activity of PdaA and its orthologs (InterPro IPR014235) is unique within the broader deacetylase family. SwsB and its orthologs (IPR014228) are distinguished from PdaA and other CE4 enzymes by two features that are correlated. SwsB contains an additional *N*-terminal domain that PdaA lacks (Figure 1c). Further, a conserved active site aspartate, D73, proposed to deprotonate water in the deacetylation step, is substituted with asparagine (Figure 1d, S2). Correspondingly, no evidence of SwsB deacetylase activity has been detected.^16^ The reaction SwsB catalyzes, and why it is essential for CwlJ-dependent germination, remains unclear. In a recent study, we identified a *B. subtilis* PdaA (*Bs*PdaA) mutant that provides a clue towards the function of SwsB.^12^ *Bs*PdaA^D73N^ cannot deacetylate MurNAc but remains able to cyclize MurN to muramic-δ-lactam.^12^ Because *Bs*PdaA^D73N^ contains the same D to N substitution that is native to wild-type SwsB, we hypothesized that SwsB is also a muramic-δ-lactam cyclase.

We purified *B. subtilis* SwsB (*Bs*SwsB) to homogeneity (Figure S3) and used a liquid chromatography-mass spectrometry (LC-MS) assay to assess muramic-δ-lactam synthesis (Figure 2a). We prepared a native cortex peptidoglycan substrate by first polymerizing Lipid II extracted from *B. subtilis* using SgtB, a glycosyltransferase from *Staphylococcus aureus*.^21,22^ The linear polymer product was incubated with *B. subtilis* CwlD (*Bs*CwlD), which cleaves the pentapeptide of every other MurNAc, followed by PdaA1 from *Clostridioides difficile* (*Cd*PdaA1), a variant that catalyzes rapid MurNAc deacetylation but slow MurN cyclization.^12^ The His_6_-tagged enzymes were then removed by passing the reaction mixture over a plug of Ni-resin to afford a linear peptidoglycan substrate enriched in MurN (“MurN-polymer”). Polymer enriched in peptide-cleaved MurNAc (“MurNAc-polymer”) was prepared by omitting addition of *Cd*PdaA1. MurNAc deacetylase and muramic-δ-lactam cyclase activities were assessed by re-addition of an enzyme to the MurNAc- or MurN-polymer substrates, respectively. After incubation, the polymer was digested with the muramidase mutanolysin to give di- or tetrasaccharide products **A**-**E** that can be detected by LC-MS (Figure 2a,b, S4). Product **E** results from borohydride reduction of lactam-containing **D** in our assay workflow, and we therefore interpret the presence of **E** as evidence of muramic-δ-lactam formation. Re-addition of *Bs*PdaA to both MurNAc- and MurN-polymer resulted in muramic-δ-lactam synthesis, as observed previously (Figure 2c,d).^12^ When *Bs*SwsB was re-added to MurNAc-polymer, only starting material **B** was detected, indicating that *Bs*SwsB is not a MurNAc deacetylase (Figure 2c). This result is consistent with the absence of a suitable catalytic base in *Bs*SwsB. However, when *Bs*SwsB was re-added to MurN-polymer, we observed complete conversion of MurN to muramic-δ-lactam; **E** was detected as the only product (Figure 2d). Thus, SwsB acts as a monofunctional muramic-δ-lactam cyclase.

**Figure 2.**
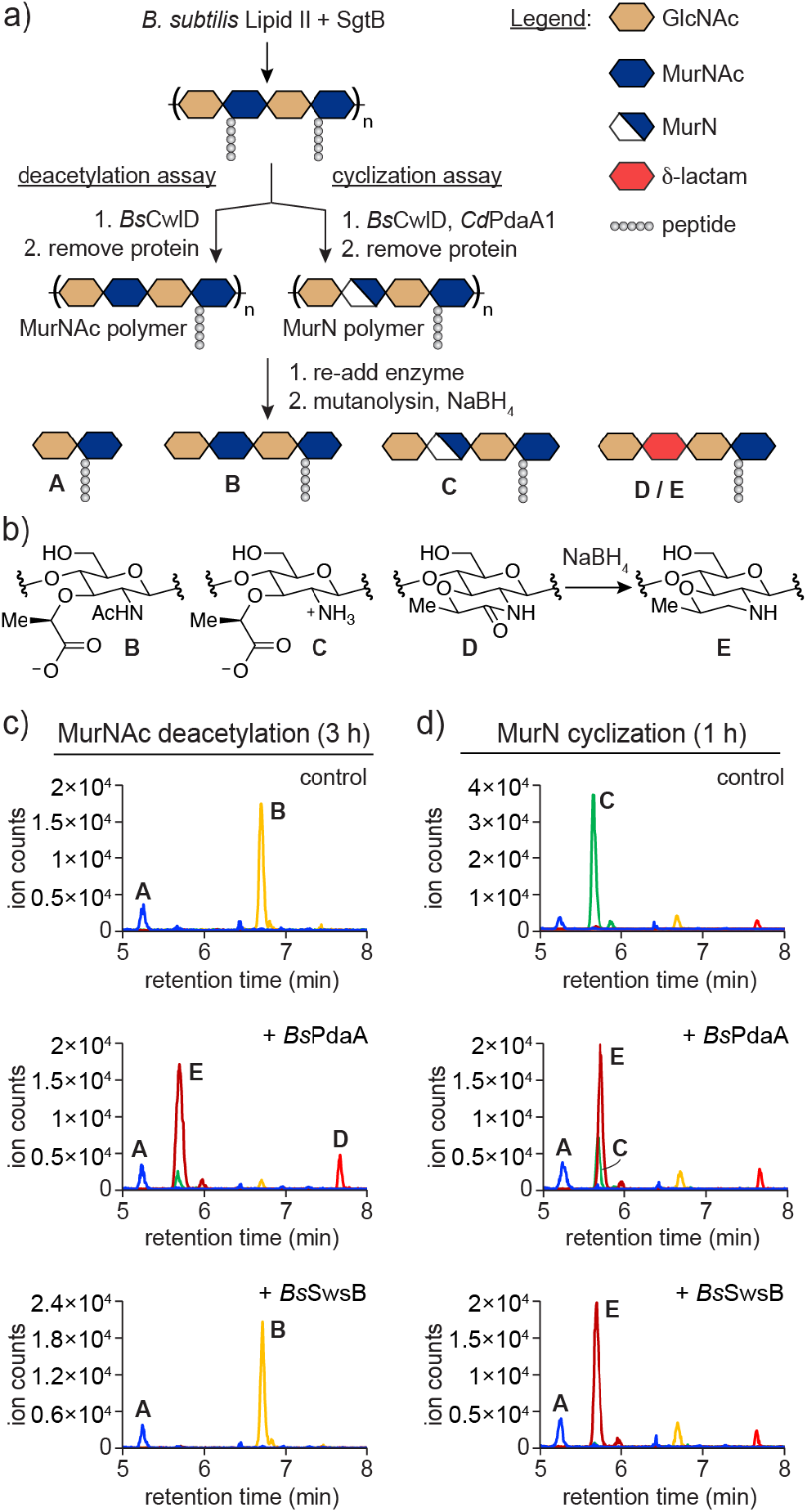
*Bs*SwsB is a muramic-δ-lactam cyclase. (a) Schematic of the LC-MS assay for MurNAc deacetylation and MurN cyclization. (b) Structures of the modified MurN residues in products **B**-**E**. LC-MS extracted ion chromatograms of *Bs*PdaA and *Bs*SwsB (c) deacetylation and (d) cyclization reactions. Enzymes were incubated with polymer substrates in pH 7.5 buffer at room temperature. Data are representative of three independent experiments.

Having successfully reconstituted SwsB activity, we sought to further characterize the enzyme. To probe the substrate preferences of *Bs*SwsB, we prepared MurN-polymer as above and digested it with mutanolysin to give **C** (Figure 2a,b). This tetrasaccharide contains only one MurN residue and therefore represents a minimal, single-turnover substrate. When *Bs*SwsB was incubated with **C** we observed complete conversion of MurN to muramic-δ-lactam (Figure S5). Analogous reactions with *Bs*PdaA gave only starting material. The two enzymes therefore have distinct substrate preferences; *Bs*SwsB will accept a short substrate while *Bs*PdaA requires longer oligosaccharides. This knowledge proved useful in designing experiments to assess the relative rate of lactam formation. We could not perform timecourse experiments on MurN-polymer because the product of mutanolysin-digestion remains a substrate for *Bs*SwsB. To determine activity over time, we incubated *Bs*SwsB with pre-digested **C**. We found that most MurN was cyclized to lactam within 1 h, and full conversion was reached by 4 h (Figures 3a, S6). Reactions with *Bs*PdaA using MurN-polymer as the substrate occurred more slowly, with reaction times of over 8 h required for complete lactam synthesis (Figure 3a).^12^

**Figure 3.**
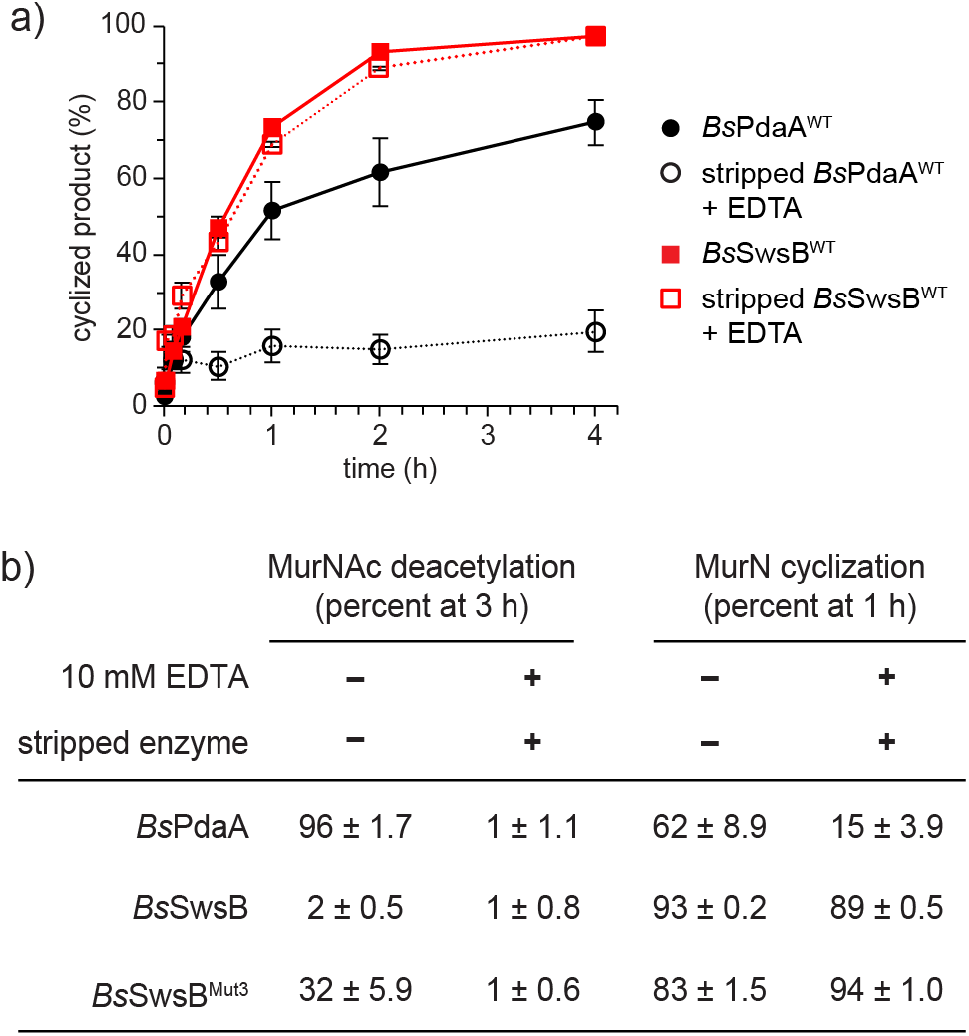
SwsB does not require a metal cofactor for cyclase activity. (a) Muramic-δ-lactam synthesis over time. Tetra-saccharide C (10 μM) and MurN-polymer were used as substrates for *Bs*SwsB and *Bs*PdaA (2 μM), respectively. Metal-stripped enzymes were incubated in buffer with 10 mM EDTA. Error bars represent the standard error of three independent experiments. (b) Percent deacetylated or cyclized products produced by *Bs*PdaA and *Bs*SwsB variants incubated with or without EDTA. Metal-stripped proteins were used for the “+ EDTA” condition. The mean and standard error of three independent experiments are reported.

The reaction catalyzed by *Bs*SwsB is notable in that an amide bond appears to form directly from a carboxylate and primary amine. Such a reaction would be energetically unfavorable, but no chemical energy was required to accomplish the transformation. We did not provide ATP in our reactions, and *Bs*SwsB has no conserved binding site for a nucleotide cofactor. Further, no obvious nucleophile is present that could serve to form an activated covalent intermediate from the substrate carboxylate. *Bs*SwsB does, however, contain a conserved metal ion-binding motif in its active site (Figure 1d). A Lewis-acid metal cofactor could serve to coordinate the MurN carboxylate and activate it for attack by the 2’-amino group. We therefore hypothesized that a metal would be required for the cyclization reaction.

To test metal dependence, we purified *Bs*SwsB and *Bs*PdaA in buffer with the chelator ethylenediaminetetraacetic acid (EDTA) to strip metal cofactors from the enzymes. We then monitored lactam synthesis over time with excess EDTA present in the reaction mixture. Reactions with stripped *Bs*PdaA were slowed in the presence of EDTA, but to our surprise, stripped *Bs*SwsB maintained robust cyclase activity, with no apparent rate decrease relative to EDTA-free conditions (Figures 3a, S6). We were concerned that these results were due to the presence of trace metal contaminants that could bind to the enzymes in an equilibrium process. We therefore evaluated MurNAc deacetylation, which is metal-dependent, under the same conditions. Stripped *Bs*PdaA had no detectable deacetylase activity, indicating the absence of a metal ion in the active site (Figure 3b). Deacetylase activity was recovered in the stripped protein by addition of excess Zn^2+^ (Figure S7). *Bs*SwsB does not catalyze deacetylation, so we could not rule out that metals were available to this enzyme. We therefore mutated *Bs*SwsB to make it a deacetylase. Three amino acid substitutions were required: N137D, A226R, and L135T. The N137D substitution re-introduces a catalytic base residue while A226R and L135T restore hydrogen bonds that may correctly position the aspartate (Figure 1d). The resulting triple mutant, *Bs*SwsB^Mut3^, showed moderate deacetylase activity when the reaction was supplemented with Zn^2+^ but deacetylation was fully quenched by EDTA (Figures 3b, S7, S8). Our reaction conditions are therefore sufficient to remove metals from the enzyme. Remarkably, metal-stripped *Bs*SwsB^Mut3^ retained full cyclase activity even in the presence of excess EDTA (Figure 3b, S6). The reaction buffer in these experiments contained 2 mM Ca^2+^, which is optimal for chemoenzymatic synthesis of the substrates.^12^ To confirm that Ca^2+^ ions were not involved, we scaled up the procedure to produce tetrasaccharide **C** and HPLC-purified the product. When purified **C** was incubated with *Bs*SwsB in buffer containing EDTA and no Ca^2+^, **E** was detected as the only product (Figure S9). From these results, we conclude that *Bs*SwsB does not require a metal ion cofactor to accomplish muramic-δ-lactam synthesis.

We have shown that *Bs*SwsB is a cyclase enzyme, synthesizing muramic-δ-lactam directly from MurN. This activity is highly unusual. Amide bond forming reactions in biology are normally accomplished by first activating the carboxylate for acyl substitution, typically as an acyl-adenylate that is transferred to an intermediate ester or thioester prior to attack by an amine nucleophile.^23^ *Bs*SwsB appears to form an amide bond directly from a carboxylic acid and primary amine with no requirement for chemical energy input. PdaA is the only other enzyme reported with similar activity,^12^ but it also acts as a deacetylase. *Bs*SwsB is unprecedented in that it has evolved to exclusively catalyze direct lactam formation.

Additional research is needed to determine the reaction mechanism, and we expect that the methods and substrates reported here will inform and enable future structural and kinetic studies. In principle, direct ligation of a carboxylate and primary amine could be accomplished by binding MurN into a reactive conformation and charge neutralization of the reacting groups by desolvation. A metal-free mechanism would necessitate re-protonation of a leaving water molecule. H222 of *Bs*PdaA is proposed to serve as a general acid during MurNAc deacetylation.^20^ Mutation of H222 in *Bs*PdaA and the corresponding H282 residue in *Bs*SwsB abolished cyclase activity (Figures 1d, S10). Therefore, it is plausible that these invariant histidines act as general and/or Lewis acids during lactam synthesis (Figures 4a, S2).

**Figure 4.**
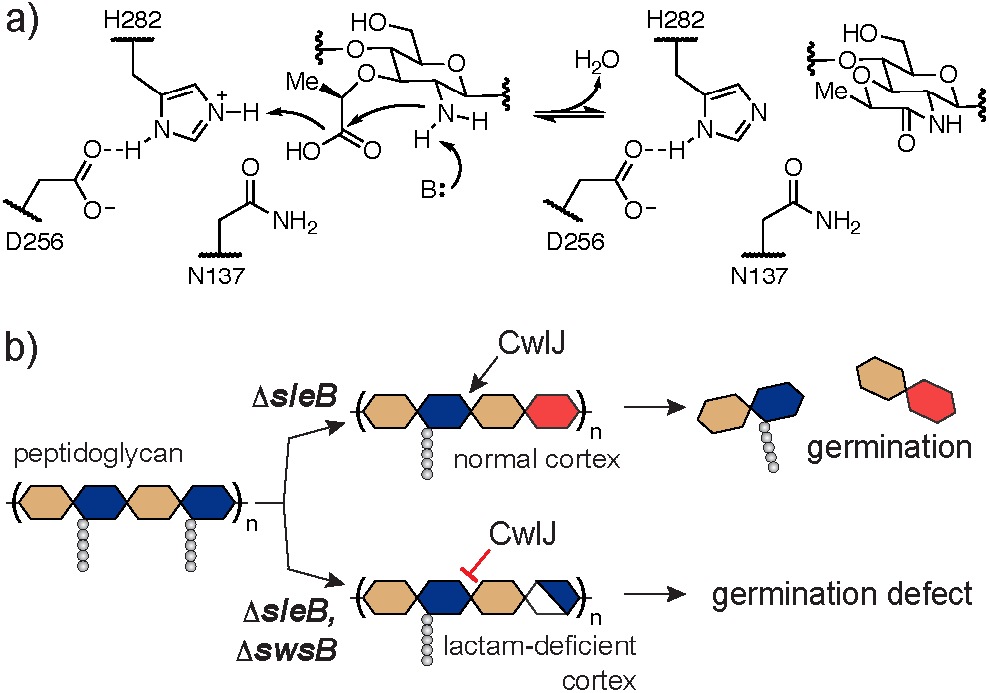
A possible mechanism for SwsB-catalyzed muramic-δ-lactam synthesis (a), and a model for the role of SwsB in CwlJ-dependent germination (b).

In the broader context of the spore, our finding that *Bs*SwsB synthesizes muramic-δ-lactam helps explain why it is essential for CwlJ-dependent germination.^16^ CwlJ is proposed to cleave cortex peptidoglycan only when muramic-δ-lactam is present.^4,16^ *Bs*PdaA catalyzes MurNAc deacetylation faster than it does MurN cyclization;^12^ therefore, the role of *Bs*SwsB may be to cyclize any MurN residues left behind by *Bs*PdaA in the outer cortex. Insufficient muramic-δ-lactam formation when *Bs*SwsB is deleted may produce a cortex peptidoglycan layer that CwlJ cannot digest, preventing germination and outgrowth (Figure 4b).

## AUTHOR INFORMATION

## Author Contributions

## ACKNOWLEDGMENT

This research was supported by Research Corporation for Science Advancement Cottrell Scholar Award CS-CSA-2024-045, Hamilton College, and the Edward and Virginia Taylor Fund for Student Research in Chemistry.

