## Supplementary Information for "Direct Amide Bond Formation By A Spore Peptidoglycan Biosynthetic Enzyme"

**METHODS**

**Materials.** Unless indicated otherwise, all reagents and chemicals were purchased from Sigma-Aldrich and used without further purification. Oligonucleotide PCR primers were purchased from Integrated DNA Technologies.
*B. subtilis* Lipid II was extracted and purified as reported.[1] *S. aureus* SgtB, *B. subtilis* CwlD, and PdaA variants (Table S1) were purified as previously described.[2,3] Mini-PROTEAN TGX gels (4-20%, Bio-Rad) were used for SDS-PAGE.

**Bacterial Culture.** The bacterial strains and plasmids used in this study are listed in Table S2. *E. coli* strains were grown at 37 °C with shaking in LB broth (Miller, Beckton Dickinson) or on agarized plates. Antibiotics were used at the following concentrations: ampicillin, 100 µg/mL; carbenicillin, 100 µg/mL; kanamycin, 50 µg/mL.

**Cloning of *B. subtilis* SwsB.** Plasmid pET24b was linearized by PCR using primers oMW47 and oMW104 (Table S3). The *swsB[K45-K319]* gene, which truncates a predicted N-terminal signal peptide and nonpolar alpha helix, was amplified from *B. subtilis* PY79 genomic DNA using the primers oMW181 and oMW182. The DNA products were gel purified and joined by isothermal assembly using Gibson Assembly Master Mix (New England Biolabs) to produce plasmid pMW1295, which expresses *B. subtilis* SwsB[K45-K319] with a C-terminal $His_6$-tag.

**Site-directed mutagenesis of *B. subtilis* SwsB and PdaA.** Site-directed mutagenesis of SwsB and PdaA was conducted by amplifying the respective expression vectors with PCR primers encoding the mutation and circularizing the linear product using kinase, ligase, Dpn1 reaction mix (KLD, New England Biolabs). Primers used for each mutation are listed in Table S3.

**Protein expression and purification – general protocol used for all proteins.** *E. coli* C43(DE3) containing the appropriate plasmid was grown in 500 mL LB broth supplemented with kanamycin at 37 ºC with shaking until the OD600 was 0.5-0.6. The culture was cooled to 16 ºC before inducing protein expression with 500 µM isopropyl-β-D-thiolgalactopyranoside (IPTG) and shaking for 16 h. Cells were harvested by centrifugation (7,000 × g, 15 min, 4 ºC) and resuspended in 20 mL lysis buffer (50 mM HEPES pH 7.5, 400 mM NaCl). The cell suspension was supplemented with DNase (0.2 mg/mL) and phenylmethanesulfonyl fluoride (1 mM) and the cells lysed by passage through a cell homogenizer (EmulsiFlex-C5, Avestin) at ≥ 10,000 psi. Cell debris was pelleted by centrifugation (20,000 × *g*, 30 min, 4 ºC). The resulting supernatant was supplemented with 20 mM imidazole and then rocked with 0.5 mL Ni-NTA resin (Qiagen) for 45 min at 4 ºC. The resin was collected in a column by gravity flow and then washed twice with 5 mL wash buffer (50 mM HEPES pH 7.5, 400 mM NaCl, 40 mM imidazole). The protein was eluted in 10 mL elution buffer (50 mM HEPES pH 7.5, 400 mM NaCl, 200 mM imidazole) and concentrated by centrifugal filtration. The protein was then further purified by fast protein liquid chromatography (FPLC, AKTA Go, Cytiva) on a Superdex 200 Increase 10/300 column (Cytiva) in a running buffer consisting of 50 mM HEPES pH 7.5, 400 mM NaCl. Pooled elution fractions were concentrated by centrifugal filtration and the protein absorbance measured at 280 nm. The predicted extinction coefficient of the protein (via ProtParam[4]) was used to estimate concentration. Proteins were diluted to 200 µM in running buffer with 10% glycerol (v/v), aliquoted, and stored at -80 ºC.

**Purification of Metal-stripped Proteins.** Proteins were expressed in *E. coli* C43(DE3) and purified by Ni-affinity pulldown at described above. The eluted protein was concentrated to 0.5-1 mL and dialyzed overnight in 1 L of 20 mM HEPES pH 7.5, 400 mM NaCl, 10 mM ethlylenediaminetetraacetic acid (EDTA). The protein was then further purified by FPLC as above in a running buffer of 50 mM HEPES pH 7.5, 400 mM NaCl, 10 mM EDTA. Proteins were diluted to 200 μM in this running buffer with 10% glycerol (v/v), aliquoted, and stored at -80 ºC.

**Biochemical reactions – general conditions.** Reaction conditions were adapted from a previously published protocol.[3] *B. subtilis* Lipid II was polymerized with SgtB, a monofunctional peptidoglycan glycosyltransferase from *Staphylococcus aureus*. Pooled polymerization reactions of up to 1 mL total volume were assembled under the following conditions: 50 mM HEPES, pH 7.5, 2 mM $CaCl_2$, 20 μM *B. subtilis* Lipid II, 0.2 μM SgtB, 10% DMSO (v/v). Reactions were incubated at room temperature for 30 min. *Bs*CwlD was added at 2 μM and the mixture incubated 1 h at room temperature, generating products enriched in peptide-cleaved MurNAc. When required, *Cd*PdaA was added at 2 μM for 10 min to generate polymer enriched in muramic acid (MurN) residues. To remove the enzymes, the entire reaction volume was passed over a 100 μL plug of settled Ni-NTA resin (Qiagen) by gravity flow twice. The eluate was then split into 50 μL aliquots. *Bs*PdaA or *Bs*SwsB were re-added at 2 μM and the reactions incubated at room temperature. Reactions were quenched by addition of 250 μL of methanol. Samples were then dried in a centrifugal evaporator and the residue resuspended in 50 μL of water.

Polymer reaction products were digested with mutanolysin from *Streptomyces globisporus* (Sigma Aldrich) to before LC-MS analysis. To 50 μL aliquots of reaction products, 8 U of mutanolysin was added and the reaction incubated at 37 ºC for 2 h with shaking. Aqueous sodium borohydride (10 mg/mL, 50 μL) was added and the reaction incubated for 30 min at room temperature. The solution pH was adjusted to ~4 by addition of 20% phosphoric acid (approximately 5 μL) and the reactions lyophilized to dryness overnight. The residue was dissolved in 25 μL of water and analyzed by LC-MS.

**LC-MS Analysis of Reaction Products.** LC-MS was conducted on a Thermo Scientific Vanquish HPLC in line with a Thermo LTQ XL ion trap mass spectrometer using electrospray ionization (spray voltage, 4500 V; spray temp, 300 ºC; capillary voltage, 46 V; capillary temp, 275 ºC) and operating in positive ion mode. Reaction products were separated on a Waters Cortecs T3 column (120 Å, 1.6 μm, 2.1 x 50 mm) equipped with a matching column guard using the following method: 0.6 mL/min eluent A (water/0.1% formic acid) for 2 min followed by a linear gradient of 0 to 17.5% eluent B (acetonitrile/0.1% formic acid) over 13 min. The peptidoglycan fragments were observed to elute between 5 and 8 min. Masses of the predicted $[M+H]^{+}$, $[M+2H]^{2+}$, and $[M+3H]^{3+}$ ions for the expected muropeptide fragments were extracted from the resulting total ion chromatograms to generate the extracted ion chromatograms displayed in the text and supplemental figures. Relative product amounts within a sample were calculated by integrating the product peaks and dividing by the total peak area. Mass spectrometry data was analyzed using Thermo FreeStyle 1.5.

**Characterization of Reaction Products.** Digested peptidoglycan products A-E were reported by our group in a prior publication and were characterized by MS/MS fragmentation as previously described.[3] Representative mass spectra of each species from LC-MS experiments are given in Figure S4.

**Cyclization Timecourse Reactions.** To assess muramic-δ-lactam cyclization over time, an 800 μL pool of peptidoglycan enriched in MurN was produced as described in the general procedure above. For *Bs*SwsB$^{WT}$ reactions, material eluting off of the Ni-NTA column was digested with mutanolysin (32 U) at 37 ºC for 2 h. For *Bs*PdaA and *Bs*SwsB$^{Mut3}$ the mutanolysin digestion was omitted. The reaction mixture was split into two 400 μL volumes and EDTA (10 mM) added to one aliquot. A *Bs*SwsB or *Bs*PdaA variant was then added at 2 μM and the reaction incubated static at room temperature. At timepoints, a 50 μL aliquot was removed and quenched by addition of 250 μL methanol. Samples were stored at -20 ºC prior to analysis. Methanolic samples were dried in a centrifugal evaporator and the residue resuspended in 50 μL of water. Mutanolysin digestion and LC-MS analysis were conducted as described above.

**Purification of tetrasaccharide C.** Tetrasaccharide C (see main text Figure 2) was obtained by scaling up our procedure for generating MurN-enriched polymer. *B. subtilis* Lipid II was extracted from 3 L of culture as reported,[1,3] and the entire purified Lipid II extract (approximately 1 μmol) reconstituted in 200 μL DMSO. A polymerization reaction of 2 mL total volume was assembled with the following conditions: 50 mM HEPES pH 7.5, 500 μM Lipid II, 2 mM $CaCl_2$, 0.2 μM SgtB, 10% DMSO (v/v). *Bs*CwlD (2 μM) was added and the mixture incubated at room temperature for 2 hours. *Cd*PdaA1 (2 μM) was then added and the reaction incubated at room temperature for 30 min. The reaction was quenched by addition of EDTA (10 mM). The resulting MurN-enriched polymer was digested with mutanolysin (40 U) for 2 hours at 37 ºC with shaking. The tetrasaccharide C product was then purified by HPLC.

HPLC was conducted on an Agilent LC system consisting of a 1290 Infinity II binary pump, a 1260 Infinity II diode array detector, and a 1260 Infinity II fraction collector. Products were separated on a Waters XBridge BEH Shield RP18 column (130 Å, 5 μm, 10 x 150 mm) using the following method: 5 mL/min eluent A (water/0.1% formic acid) for 3 min followed by a linear gradient of 0 to 20% eluent B (acetonitrile/0.1% formic acid) over 30 min. The two tetrasaccharide C anomers were observed to elute between 10 and 12 mins with residual product A partially co-eluting. Fractions containing the product were pooled and lyophilized to dryness. The resulting residue was reconstituted in 200 μL water and quantified by derivatization with fluorescamine in 400 mM borate pH 9.7 using L-alaninamide as a standard. Estimated yields for this procedure were low (~5 %), but enough product was obtained for ~20 small scale reactions, as below.

**Cyclization Reactions with Purified Tetrasaccharide C.** Reactions with purified tetrasaccharide C were assembled under the following conditions: 50 mM HEPES, pH 7.5, 20 μM tetrasaccharide C, 2 μM *Bs*PdaA or *Bs*SwsB variant, 10% DMSO (v/v). Reactions were incubated at room temperature for 1 h. Aqueous sodium borohydride (10 mg/mL,

50 μL) was added and the mixture incubated 30 min at room temperature. The solution pH was adjusted to ~4 by addition of 20% phosphoric acid and lyophilized to dryness. Products were analyzed by LC-MS as above.

### SUPPLEMENTAL FIGURES AND TABLES

**Table S1: UniProt accession codes for proteins.**

| Protein | Accession |
|---|---|
| *Staphylococcus aureus* SgtB† | Q93Q23 |
| *Bacillus subtilis* CwlD | P50864 |
| *Bacillus subtilis* PdaA | O34928 |
| *Bacillus subtilis* SwsB* | P50850 |
| *Clostridioides difficile* PdaA1 | Q18BV2 |

† SgtB is also known as Mgt1
* SwsB was previously known as YlxY

**Table S2: Bacterial strains and plasmids**

| Strain or plasmid | Description[a] | Reference |
|---|---|---|
| *E. coli* | | |
| C43(DE3) | BL21(DE3) derivative for protein expression | 5 |
| *Plasmids* | | |
| pET24b | IPTG-inducible protein expression vector; $Kan^R$ | Novagen |
| pET28b(+) | IPTG-inducible protein expression vector; $Kan^R$ | Novagen |
| pMgt1† | *S. aureus* SgtB-$His_6$ expression vector; $Amp^R$ | 6 |
| pMW1008 | *B. subtilis* $His_6$-CwlD[N27-E237] expression vector, $Kan^R$ | 3 |
| pMW1016 | *C. difficile* $His_6$-PdaA[S29-K242] expression vector, $Kan^R$ | 3 |
| pMW1070 | *B. subtilis* PdaA[V24-L263]-$His_6$ expression vector, $Kan^R$ | 3 |
| pMW1159-6 | *B. subtilis* PdaA[V24-L263]$^{D73N}$-$His_6$ expression vector, $Kan^R$ | 3 |
| pMW1159-9 | *B. subtilis* PdaA[V24-L263]$^{H222A}$-$His_6$ expression vector, $Kan^R$ | This study |
| pMW1295 | *B. subtilis* SwsB[K45-K319]-$His_6$ expression vector, $Kan^R$ | This study |
| pMW1304 | *B. subtilis* SwsB[K45-K319]$^{N137D}$-$His_6$ expression vector, $Kan^R$ | This study |
| pMW2022 | *B. subtilis* SwsB[K45-K319]$^{N137D,A226R}$-$His_6$ expression vector, $Kan^R$ | This study |
| pMW2024 | *B. subtilis* SwsB[K45-K319]$^{L135T,N137D,A226R}$-$His_6$ expression vector, $Kan^R$ | This study |
| pMW2067 | *B. subtilis* SwsB[K45-K319]$^{H282A}$-$His_6$ expression vector, $Kan^R$ | This study |

[a] Abbreviations: $Amp^R$, ampicillin resistance; $Kan^R$, kanamycin resistance
† SgtB is also known as Mgt1.

**Table S3: Oligonucleotide PCR primers**

| primer | Sequence (5'-3') | Mutation |
|---|---|---|
| oMW47 | CTCGAGCACCACCACCAC | |
| oMW104 | CATATGTATATCTCCTTCTTAAAGTTAAACAAAATTATTTCTAGAGG | |
| oMW129 | TTACCTGCTTGCGACCGTATCGAGGGACAATGCAGAAGCGCTGG | PdaA$^{H222A}$ |
| oMW130 | ATGGCTCCCGGGTGCGCC | PdaA$^{H222A}$ |
| oMW179 | TTTTTTAATCGATGTGGCATGGG | SwsB$^{N137D}$ |
| oMW180 | GCCACCATCGGTTTG | SwsB$^{N137D}$ |
| oMW181 | AAGAAGGAGATATACATATGAAAGACCCGTTATATGAAG | |
| oMW182 | TGGTGGTGGTGGTGCTCGAGCTTCAATAGTCTTGTTTCATC | |
| oMW192 | AAAGTGGTTCCGCCCGCCAAGCG | SwsB$^{N137D,A226R}$ |
| oMW193 | GGCTTAACGCCGATC | SwsB$^{N137D,A226R}$ |
| oMW195 | ATCGGTTTGTCAGGATTTC | SwsB$^{L135T,N137D,A226R}$ |
| oMW200 | GGTGGCTTTTACCATCGATGTGG | SwsB$^{L135T,N137D,A226R}$ |
| oMW201 | GATTTTAATGGCGCCGACTGACCCTACG | SwsB$^{H282A}$ |
| oMW202 | ATGGCACCATTATGTATC | SwsB$^{H282A}$ |

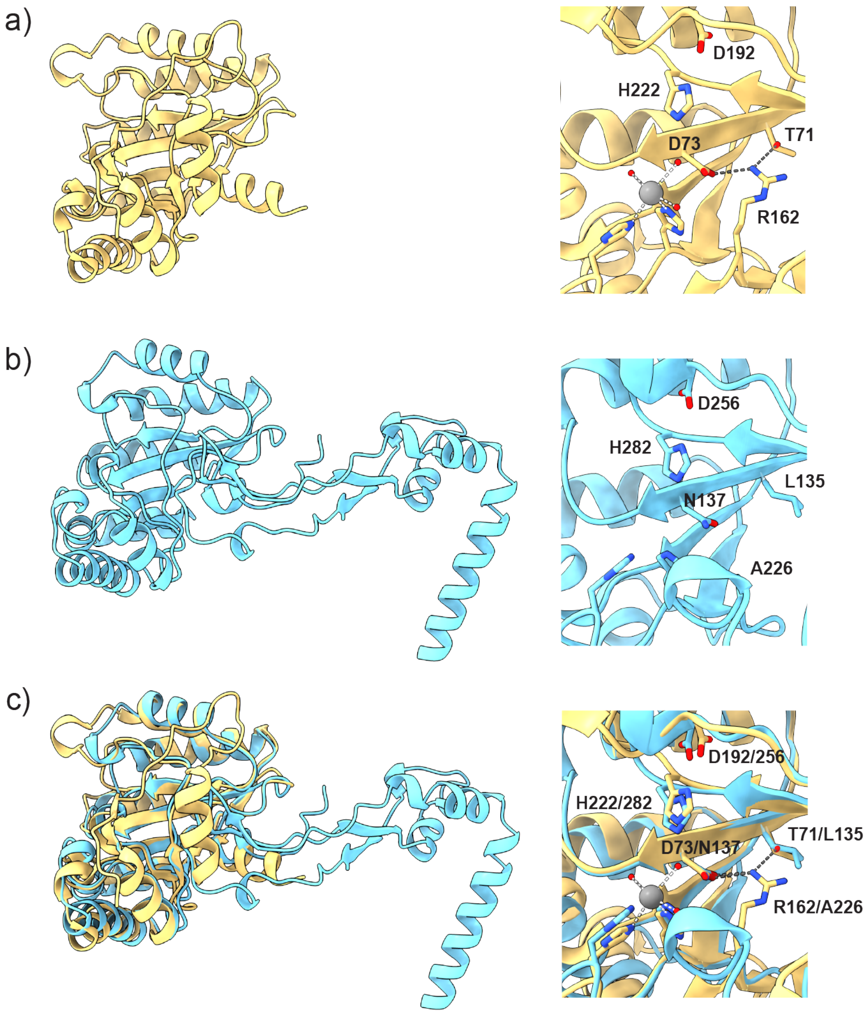


**Figure S1.** Structures of (a) *B. subtilis* PdaA (PDB 1W1B[7]) and (b) *B. subtilis* SwsB (AlphaFold3 model). The predicted *N*-terminal signal peptide is hidden in the SwsB structure. A bound divalent metal ion is shown as a gray sphere in the PdaA structure. Hydrogen bonds are shown as gray dashed lines. Coordinated water molecules are red spheres. (c) Structural alignment generated in ChimeraX version 1.9.

```
Bs PdaA         ---MKWMCSICCAAVLLAG--GAAQ---AE-----------------------------A    23
Vh PdaA         ---MRNQVKILISILLFLSI---SL---PH-----------------------------T    22
Bc PdaA         -MKYKWLYMGLIFSIM--MALVPVS---AL-----------------------------A    25
Pm PdaA         ---MRRFTFFLVFLFVMSTGIASVE---AQ-----------------------------M    25
Bs SwsB         --MYKKFVPFAVFLFLFFVSFEMMENPHALDYIGAMKKDTVTVTASKDPLYEELLQKAPE    58
Vh SwsB         MYRLRQFITFCLFLLVVGISFNLDYNPFSEREGE----QLIQTTAKRDPLYQEILQKSSD    56
Bc SwsB         --MKVRILA-----------YICIFSLYVS--------LGSYSVFAQDNLHEEIQKHAKK    39
Pm SwsB         --MKRTIVQFTAFLFLLAITYKSIYNPFAEAYIEALKSDVQLVSAQHDALYQKIEEKAKD    58

Bs PdaA         VPNEPI------NWGFKRSVNHQPPDAGKQLNSLIE-----------------------K    54
Vh PdaA         ALAGGY------GWGYKKNNNHEIPDVGK-YKDMLD-----------------------K    52
Bc PdaA         YTNTPH------NWGIPRPKNETVPDAGKLYTDLLQ-----------------------K    56
Pm PdaA         YPNTPI------SWGFQKSKNHKPASAGTAYEQILA-----------------------K    56
Bs SwsB         YEVKPQNARIDKVWKSIPGYNGLKVNIEQSYKKMKQHGKFREKDLVYSQVKPSVHLESLQ    118
Vh SwsB         YAEEAQDAYIDKVWKKTPGRNGLQVNLEKSYRNMKESGEFDENLLSLEQTSPKISLEDLP    116
Bc SwsB         YEIAPQNAMIDKIWKATPGYNGRQVDIEASYNNMKKLKEFDQKYLEFKEVSPSVHLEDLS    99
Pm SwsB         YEKPAANARIDPVWKRVPGYNGIKVDLAASYKNMKPAGKFDEKKLVYKQVRPKVHLSDLP    118

Bs PdaA         YDAFYLGNTKEKTIYLTFDNGYENGYTPKVLDVLKKHRVTGTFFVTGHFVKDQPQLIKRM    114
Vh PdaA         YGAYYADFSGEKNIYLTFDNGYEEGYTDNILDVLKKEKVPATFFVTGHYVKDQPELVKRM    112
Bc PdaA         NGGFYLGDTKKKDIYLTFDNGYENGYTGKILDVLKEKKVPATFFVTGHYIKTQKDLLLRM    116
Pm PdaA         YDAFYLGDTNKKNIYLTFDNGYENGYTPQVLDVLKKRKVPAMFFVTGHYLKEEPKLIKRM    116
Bs SwsB         PEPIYKGNPDKPMVAFLINVAWGNEYLEKMLPILQKHQVKATFFLEGNWVRNNVQLAKKI    178
Vh SwsB         SAPIYRGHPEKEMVAFLINVSWGAEYIPDILNELKKEKVKATFFIEGKWAKENAELVKMI    176
Bc SwsB         PAPIYRGHPNKKMVGLTINVAWGNEYLPRILEILKKHDVKATFFLEGRWVKENLRFAKMI    159
Pm SwsB         QEPIYRGHDEKPMVSFTVNVAWGNEYLPKMLEVLKKHHAKATFFLEGKWVKNNPDMAKMI    178

Bs PdaA         SDEGHIIGNHSFHHPDLTTKTADQIQDELDSVNEEVYKITGKQDNLYLRPPRGVFSEYVL    174
Vh PdaA         VDEGHIIGNHSYHHPDFTIMDKDKIKKELQTLEKAVAEVSDQKSLRYVRPPRGTFSENTL    172
Bc PdaA         KDEGHIIGNHSWSHPDFTAVNDEKLREELTSVTEEIKKVTGQKEVKYVRPPRGVFSERTL    176
Pm PdaA         VKEGHIVGNHSWHHPDLTQVDDARFKEELQKVKDEYKNITGRDEMKYLRSPRGVFSERTL    176
Bs SwsB         AKDGHEIGNHSYNHPDMSKLTTGRISEQLDKTNEQIEQTIGVK-PKWFAPPSGSFRKAVI    237
Vh SwsB         SEQGHVIGNHAYNHPDMARLSNQKNIEQISSTNEIIKAIIDEE-PKWFAPPSGSYNQHVV    235
Bc SwsB         VDANQEVGNHSYTHPNMKTLSSDEIREQLQKTNRMIEVVTNQK-VRWFAPPSGSFRDEVV    218
Pm SwsB         VDAGHEVGNHSYSHPDMATLSASQINQQLKKTNDIITSTTGQK-VKWFAPPSGSFRPEVV    237

Bs PdaA         KETKRLGYQTVFWSVAFVDWKINNQKGKKYAYDHMIKQAHPGAIYLLHTVSRDNAEALDD    234
Vh PdaA         KWTYDLGYTHIFWSLAFIDWHTKKQKGWKYAYEQVMDQIHPGAIVLLHTVSSDNAEALQH    232
Bc PdaA         ALTKEMGYYNVFWSLAFLDWKVDQQRGWQYAHNNVMTMIHPGSILLLHAISKDNAEALAK    236
Pm PdaA         ALSKQEGYTNVFWSLAFVDWKVNEQKGWRYSYDNMMAQIHPGAIMLLHTVSKDNADALDQ    236
Bs SwsB         DIAAEKQMGTVMWTVDTIDWQKPAP---SVLQTRVLSKIHNGAMILMHPTDP-TAESLEA    293
Vh SwsB         DAAHNLNMHTILWTVDTIDWKNPTV---SVMINRVNDKLHPGATILMHPTES-IAEGIGP    291
Bc SwsB         KIADDFQMGTIMWTVDTIDWKRPEP---DVLLQRVMRKIHPGAIVLMHPTSS-TAEALDT    274
Pm SwsB         TLASQLKMSTIMWTVDTIDWQKPSS---EVLINRVMKKIHPGAIVLMHPTES-TAESLDQ    293

Bs PdaA         AITDLKKQGYTFKSIDDLMFEKEMRLPSL----      263
Vh PdaA         MITELKKQGYSFKSLDELVMKDHIPKVIYGLE-      264
Bc PdaA         IIDDLREKGYHFKSLDDLVKGNQP---------      260
Pm PdaA         AIVDLKKQGYTFKSIDDLMNERKQMKKSPSETK      269
Bs SwsB         LITQIKDKGYALGTVTELMDETRLLK-------      319
Vh SwsB         LIRQVKQKGFKLGTIEKLLNEER----------      314
Bc SwsB         MIKKLKEQGYKVGNITELLDEKRVD--------      299
Pm SwsB         LLTDIERKGLKVSDVSTMLDEERMMKIPSSTKK      326
```

text = conserved structural or substrate-/cofactor-binding residues
text = semi-conserved or complementary structural residue
text = catalytic or active site residue, conserved in both proteins
text = catalytic or active site residue, conserved in PdaA variants
text = catalytic or active site residue, conserved in SwsB variants

**Figure S2.** Multiple protein alignment of selected PdaA and SwsB variants. Sequences were obtained from the KEGG genome database (https://www.genome.jp/kegg/genome) and aligned using NCBI Protein BLAST. Species labels are *Bs – Bacillus subtilis* 168, *Vh – Virgibacillus halodenitificans* PDB-F2, *Bc – Bacillus cereus* ATCC 14579, *Pm – Priestia megaterium* QM B1551. The UniProt entries for each protein are as follows: *Bs* PdaA – O34928, *Bs* SwsB – P50850, *Vh* PdaA – A0AAC9IXT6, *Vh* SwsB – A0AAC9J0E3, *Bc* PdaA – Q81IC4, *Bc* SwsB – Q819Z2, *Pm* PdaA – D5DXQ5, *Pm* SwsB – D5DQ55.

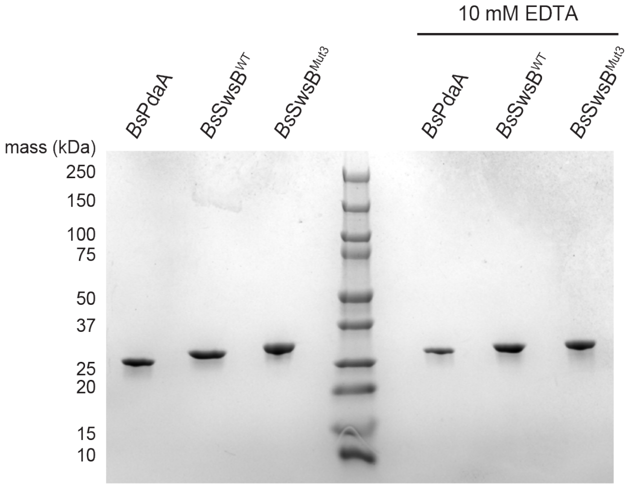


**Figure S3.** Coomassie-stained SDS-PAGE of purified proteins. Proteins in the "10 mM EDTA" lanes were purified according to the metal-stripping protocol (see Methods). Protein was loaded at approximately 2 μg per lane.

**Peak A**

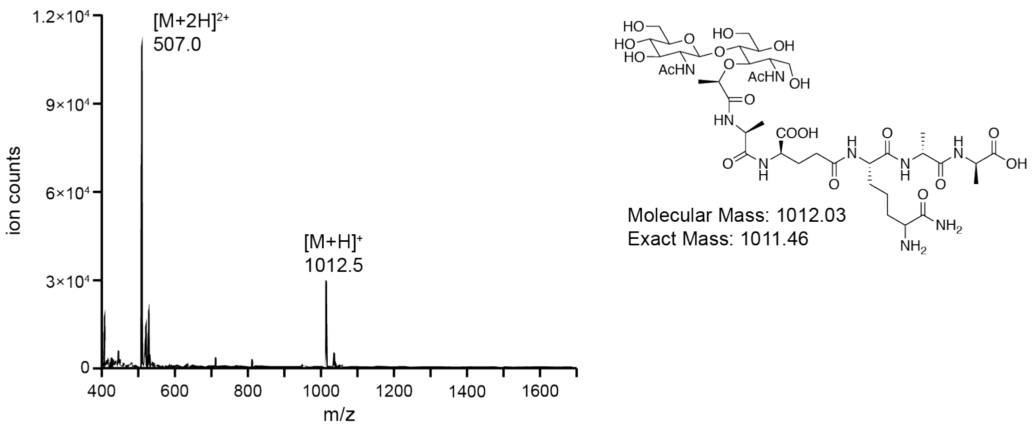


**Peak B**

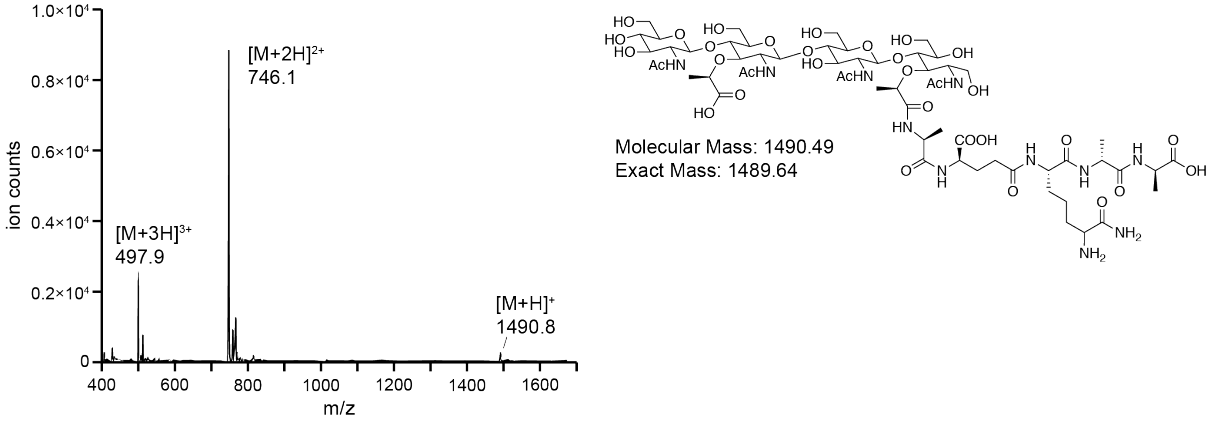


**Peak C**

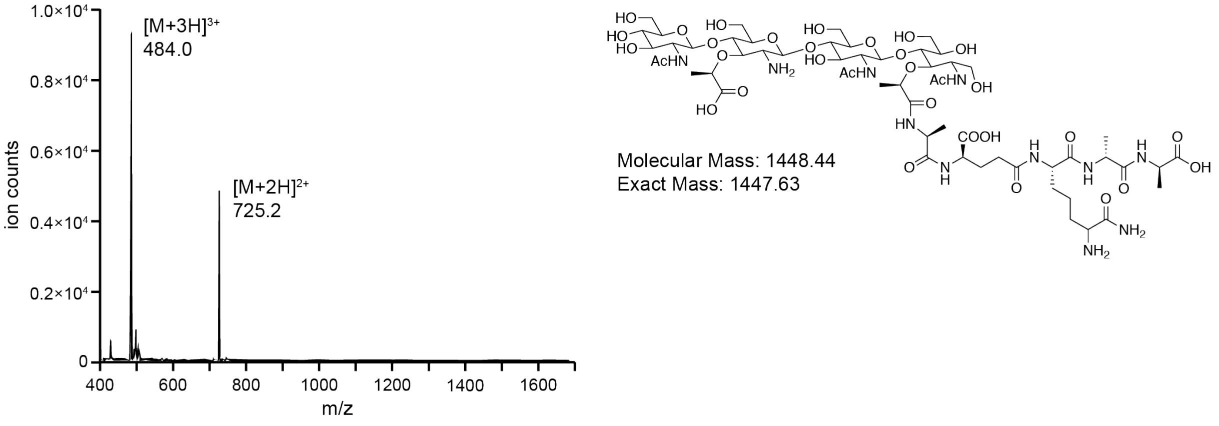


**Figure S4.** Electrospray ionization mass spectra of peptidoglycan fragments. Products A-E are defined as in the main text. The diagnostic ion adducts are labeled for each species.

**Peak D**

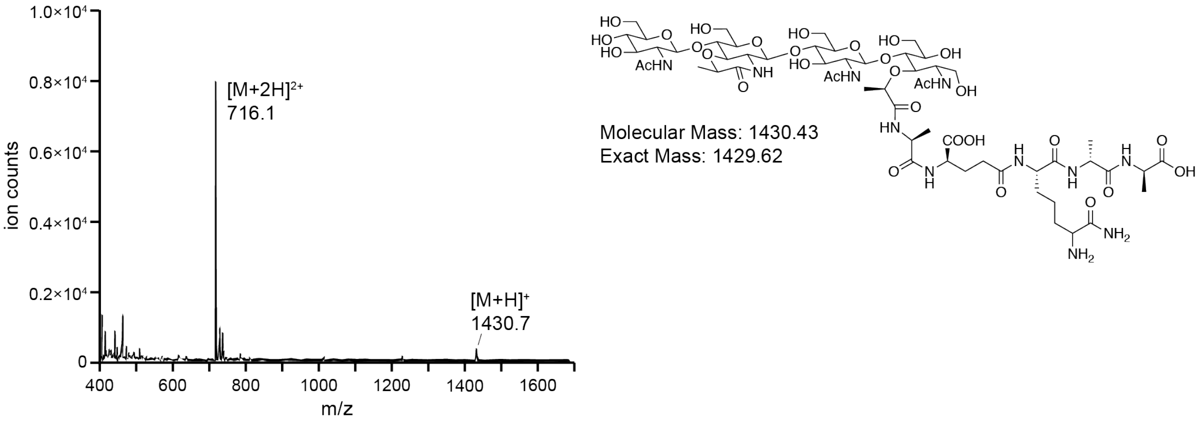


**Peak E**

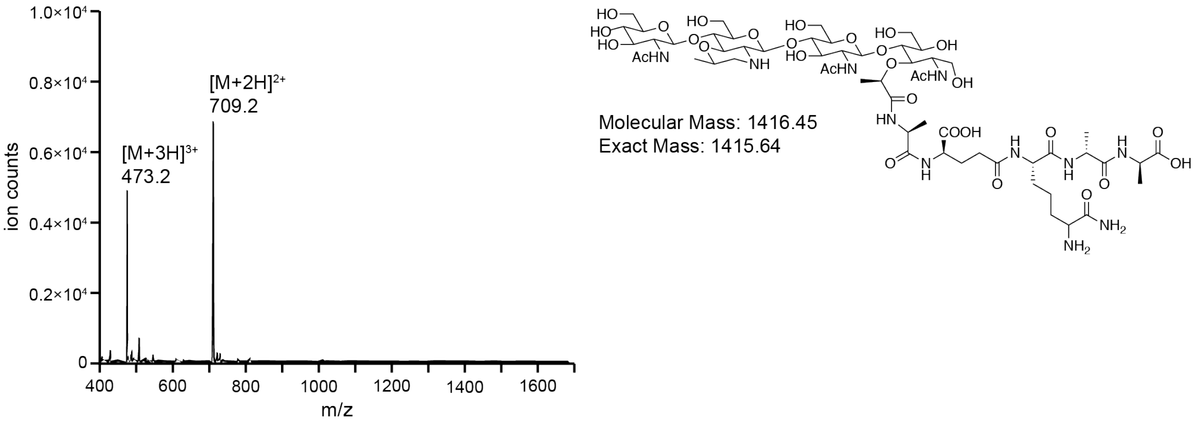


**Figure S4 (continued).** Electrospray ionization mass spectra of peptidoglycan fragments. Products A-E are defined as in the main text. The diagnostic ion adducts are labeled for each species.

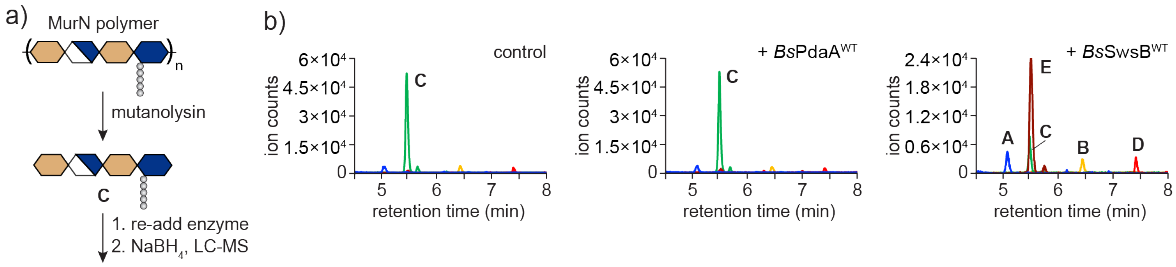


**Figure S5.** *Bs*SwsB will accept digested peptidoglycan as a substrate. a) Schematic of experimental workflow. Peptidoglycan enriched in MurN was digested with mutanolysin to produce tetrasaccharide C. Enzymes were re-added to the digested material for 1 h and the reaction products analyzed by LC-MS. b) LC-MS extracted ion chromatograms of an untreated control and reactions with *Bs*PdaA or *Bs*SwsB. Data are representative of two independent experiments.

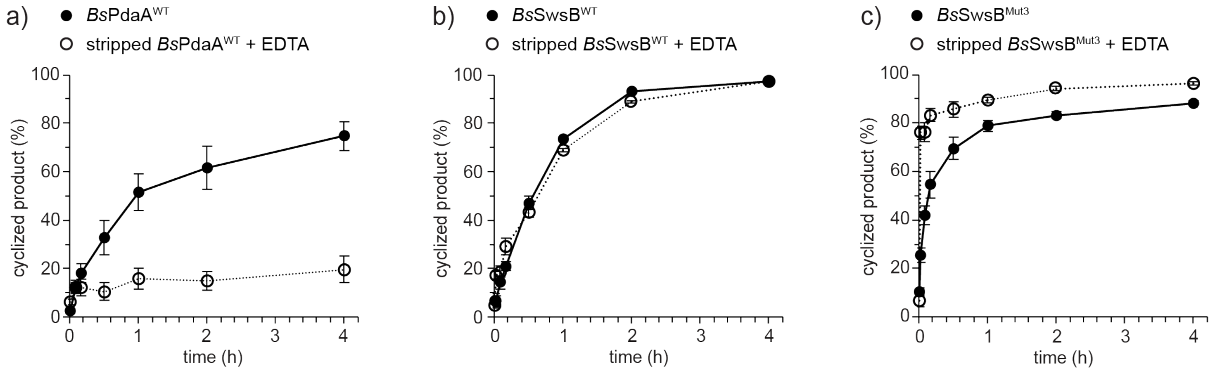


**Figure S6.** Analysis of muramic-δ-lactam synthesis over time. Pooled reactions were assembled using MurN polymer as a substrate for (a) *Bs*PdaA and (c) *Bs*SwsB$^{Mut3}$ or digested tetrasaccharide C for (b) *Bs*SwsB$^{WT}$. Open circles are reactions with metal-stripped protein (2 µM) in the presence of 10 mM EDTA. At timepoints, aliquots were removed, methanol-quenched, and the products were analyzed by LC-MS. Error bars represent the standard error of three independent experiments. Data in panels a and b is reproduced from Figure 3a.

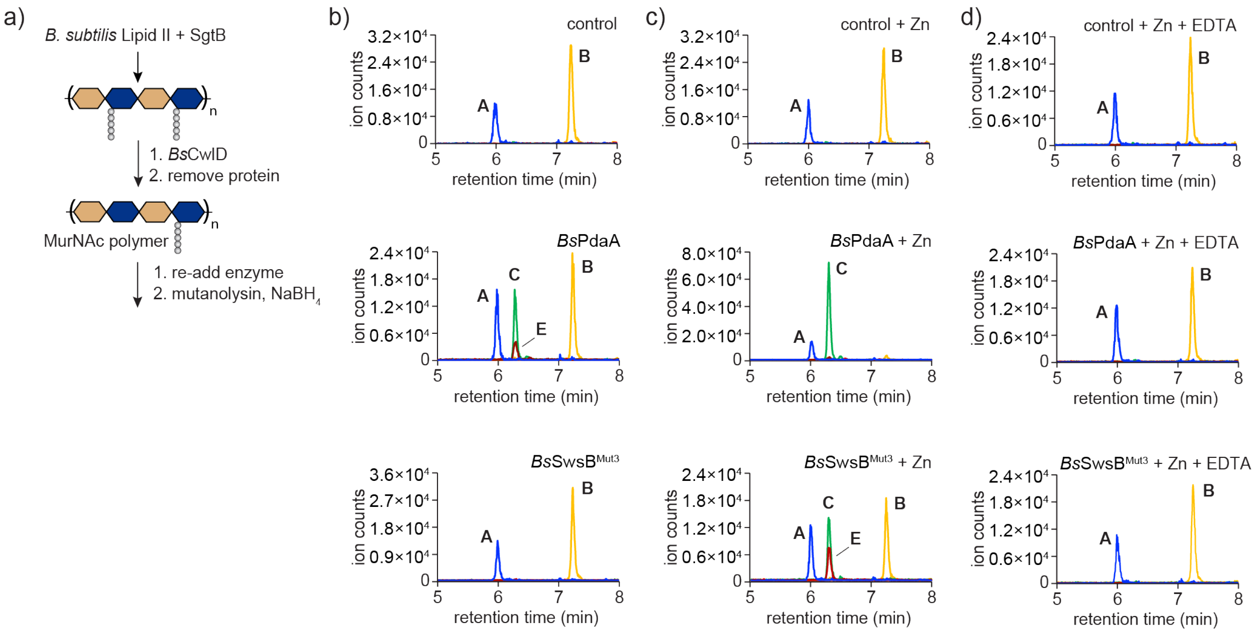


**Figure S7.** Deacetylase activity can be recovered in the metal-stripped proteins by addition of $Zn^{2+}$. (a) Schematic of experimental workflow. Linear peptidoglycan was incubated with *Bs*CwlD for 1 h to produce polymer enriched in peptide-cleaved MurNAc. Metal-stripped *Bs*PdaA or *Bs*SwsB$^{Mut3}$ (2 μM) were added to reactions supplemented with (b) no additive, (c) 110 μM $ZnCl_2$, or (d) 110 μM $ZnCl_2$ + 10 mM EDTA (d). The 110 μM $ZnCl_2$ concentration was chosen because the metal-stripped protein aliquots contain EDTA themselves (see Methods), giving a final EDTA concentration of 100 μM upon dilution into the reaction mixture. Reactions were incubated 1 h at room temperature and the products analyzed by LC-MS. The data are representative of two independent experiments.

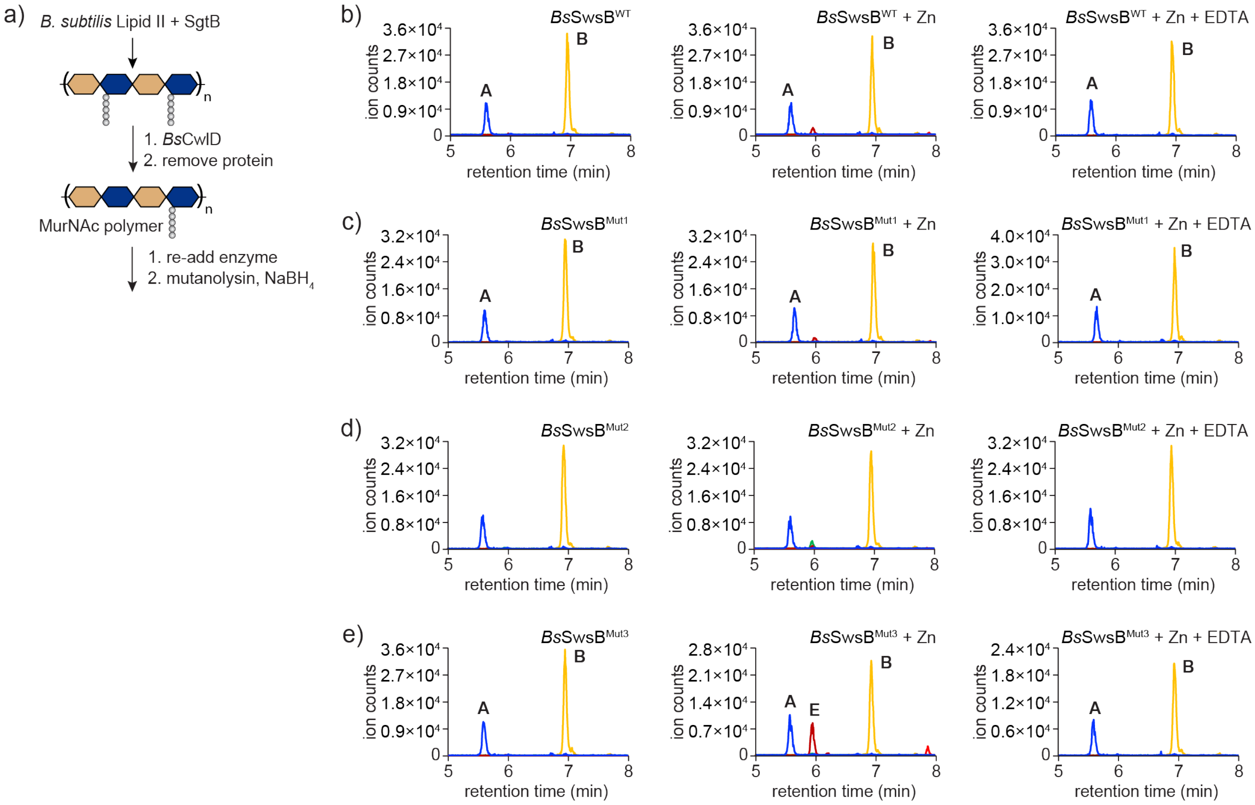


**Figure S8.** Generation of a SwsB variant with deacetylase activity. (a) Schematic of experimental workflow. Linear peptidoglycan was incubated with *Bs*CwlD for 1 h to produce polymer enriched in peptide-cleaved MurNAc. Polymer aliquots were supplemented with 2 μM $ZnCl_2$ or 2 μM $ZnCl_2$ + 5 mM EDTA. Purified SwsB variants were added at 2 μM and the reactions incubated at room temperature for 3 h. The variants are as follows: (b) wild-type *Bs*SwsB; (c) *Bs*SwsB$^{Mut1}$ = N137D; (d) *Bs*SwsB$^{Mut2}$ = N137D,A226R; (e) *Bs*SwsB$^{Mut3}$ = N137D,A226R,L135T. Data are representative of three independent experiments.

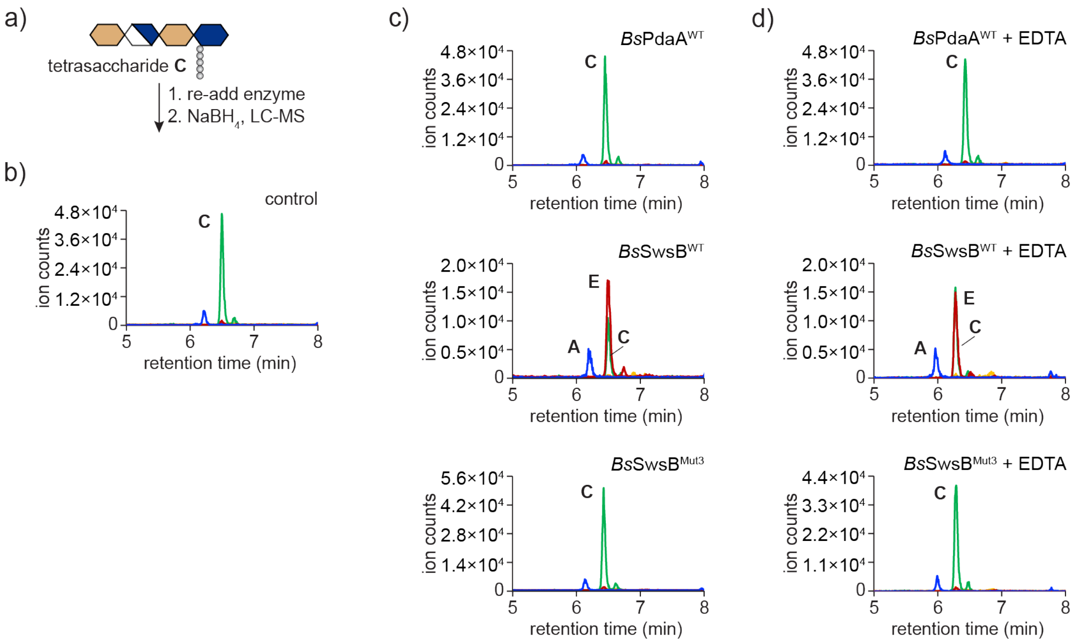


**Figure S9.** Reactions with purified tetrasaccharide C. (a) Schematic of experimental workflow. See Methods for the experimental protocol to produce purified C. Tetrasaccharide C (20 μM) and the indicated enzymes (2 μM) were incubated in 50 mM HEPES pH 7.5 at room temperature for 1 h and the products analyzed by LC-MS. (b) No enzyme control. (c) Reactions without EDTA in the reaction buffer. (d) Reactions in buffer supplemented with 10 mM EDTA. The data are representative of two independent experiments.

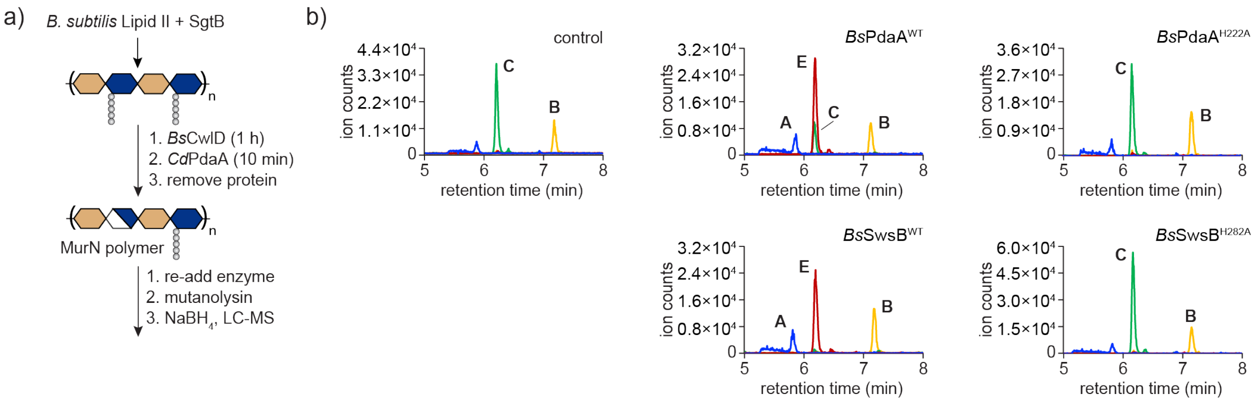


**Figure S10.** Mutation of an invariant active site His abolishes cyclase activity. (a) Schematic of experimental workflow. Enzymes (2 μM) were incubated with MurN-enriched polymer for 1 h at room temperature. (b) LC-MS extracted ion chromatograms of cyclization reaction products. The data are representative of two independent experiments.
